# Shallow Genome Sequencing for Phylogenomics of Mycorrhizal Fungi from Endangered Orchids

**DOI:** 10.1101/862763

**Authors:** Sarah A. Unruh, J. Chris Pires, Lawrence Zettler, Luigi Erba, Igor Grigoriev, Kerrie Barry, Christopher Daum, Anna Lipzen, Jason E. Stajich

**Affiliations:** Division of Biological Sciences, University of Missouri, Columbia, MO, United States; Department of Biology, Illinois College, Jacksonville, IL, United States; United States of America Department of Energy Joint Genome Institute, Walnut Creek, CA, United States; Microbiology and Plant Pathology, University of California-Riverside, Riverside, CA, United States

## Abstract

Most plant species form symbioses with mycorrhizal fungi and this relationship is especially important for orchids. Fungi in the genera *Tulasnella, Ceratobasidium,* and *Serendipita* are critically important for orchid germination, growth and development. The goals of this study are to understand the phylogenetic relationships of mycorrhizal fungi and to improve the taxonomic resources for these groups. We identified 32 fungal isolates with the internal transcribed spacer region and used shallow genome sequencing to functionally annotate these isolates. We constructed phylogenetic trees from 408 orthologous nuclear genes for 50 taxa representing 14 genera, 11 families, and five orders in Agaricomycotina. While confirming relationships among the orders Cantharellales, Sebacinales, and Auriculariales, our results suggest novel relationships between families in the Cantharellales. Consistent with previous studies, we found the genera *Ceratobasidium* and *Thanatephorus* of Cerabotasidiaceae to not be monophyletic. Within the monophyletic genus *Tulasnella*, we found strong phylogenetic signals that suggest a potentially new species and a revision of current species boundaries (e.g. *Tulasnella calospora*); however it is premature to make taxonomic revisions without further sampling and morphological descriptions. There is low resolution of *Serendipita* isolates collected. More sampling is needed from areas around the world before making evolutionary-informed changes in taxonomy. Our study adds value to an important living collection of fungi isolated from endangered orchid species, but also informs future investigations of the evolution of orchid mycorrhizal fungi.

## INTRODUCTION

Fungi are more than mere decomposers, they form symbioses with every other group of organisms on Earth. Fungal interactions span the entire symbiotic spectrum, from parasitism to mutualism. Most intertwined with plants, may have even enabled development/existence of land plants (Lutzoni et al., 2018). As a result of this long-term association, fungi are essential symbionts to almost every plant species on Earth. The fungi live in plant roots are called mycorrhizal fungi and associate with more than 85% of plant species (Smith and Read, 2008). Mycorrhizal fungi are critical for plant health and function by helping obtain and retain water, mediating defense responses, participating in signaling between roots, and facilitating the exchange of nutrients like carbon, phosphorus, and nitrogen (Barto et al., 2012; Jung et al., 2012; Peterson and Massicotte, 2004; Wang et al., 2017; Yoder et al., 2010). The plant group that relies the most on their mycorrhizal fungi are orchids.

Orchids rely on their mycorrhizal symbionts to stimulate plant development during seed germination by providing carbon resources (Kuga et al., 2014). Orchid mycorrhizal fungi (ORM) form hyphal coils termed pelotons inside the cells of orchid embryos and in the adult roots, tubers, or rhizomes (McCormick et al., 2016; Rasmussen et al., 2015). These pelotons are the sites of nutrient exchange and the molecular nature of this marketplace remains poorly understood though exciting new research shedding light (Fochi et al., 2017a; Fochi et al., 2017b; Kuga et al., 2014). Most orchids associate with mycobionts belonging to the basidiomycete groups Sebacinales, Ceratobasidiaceae and Tulasnellaceae. In addition to orchid mycorrhizal fungi, these groups contain saprotrophs, plant pathogens, and ectomycorrhizal representing a wide array of metabolic capabilities (Kohler et al., 2015; Nagy et al., 2016). Furthermore, molecular studies have revealed simultaneous root colonization by multiple fungal partners in both photosynthetic terrestrial and epiphytic orchids (Martos et al., 2012). Concluding sentence that makes the argument that there are many dynamics we need to better understand so we need to characterize the diversity of these fungi to untangle their interactions and mechanisms.

Although fungi play critical roles, they are rarely visible on the landscape. The number of extant fungal species on Earth ranges from 2-5 million (Blackwell, 2011; Hawksworth and Lücking, 2017) up to 166 million species (Larsen et al., 2017). Most species are microscopic and over the last few decades species identification has relied on molecular methods. Historically, these methods often have used a single molecular marker such at ITS (Nilsson et al., 2014). However modern genome sequencing methods are important tools to discover and describe taxonomic, phylogenetic and functional diversity. The use of different, new analytical tools has also greatly benefited our knowledge of the below-ground ecology of orchids and orchid mycorrhizal fungi. On the right track with multiple markers and Bayesian species delimitations (Ruibal et al., 2014; Ruibal et al., 2013; Whitehead et al., 2017). New species of Tulasnella and relatives are constantly being identified (Linde et al., 2017). Continue to combine sequencing with taxonomic knowledge to provide a comprehensive description of the species that associate with orchids.

The genera of orchid fungi we have sampled belong to two orders, Cantharellales and Sebacinales, in the Agaricomycetes. Cantharellales is sister to the rest of class Agaricomycetes and comprises seven families total (Ceratobasidiaceae, Tulasnellaceae, Botryobasidiaceae, Cantharellaceae, Clavulinaceae, Hydnaceae, and Aphelariaceae), though Hibbett et al., (2014), define Cantharellaceae and Clavulinaceae as synonymous with Hydnaceae and the status of Aphelariaceae is unknown (Kirk et al., 2008; Leacock, 2018). Ceratobasidiaceae has two genera (*Ceratobasidium* and *Rhizoctonia*/*Thanatephorus*) that have been demonstrated to be polyphyletic (Veldre et al., 2013). In fact, the type specimen for *Ceratobasidium* has since been reclassified as a member of the order Auriculariales based on the characters like the shape of the basidia and the dolipore (specialized hyphal septa) ultrastructure, leading Oberwinkler et al., (2013a) to restrict *Ceratobasdium* and Ceratobasidiaceae to the type specimen and reclassifying *Ceratobasidium* spp. as *Rhizoctonia* (Kirk et al., 2008). Tulasnellaceae contains 3 genera and c. 50 sp (Kirk et al., 2008). In addition to these described families, the genus *Sisotrema* is known to be polyphyletic with members in Auriculariales as well as Cantharellales. Successively sister to the rest of the Agaricomycetes is the order Sebacinales which includes two families – the Sebacinaceae and Serendipitaceae (Weiss et al., 2016). Though this order comprises a wide swath of diversity, it remains difficult to adequately describe species due to a high volume of environmental sequence data without information about morphological characters (Oberwinkler et al., 2013b; Weiss et al., 2016).

In this study, our primary goal is to shallowly sequence a rich living collection of fungi isolated from orchid roots and seedlings to provide a phylogenetic framework for future genome-enabled evolutionary and functional studies. Our secondary goal, with the addition of key outgroups, is to answer a series of nested phylogenetic questions about the relationships among the orders, families and genera of Agaricomycetes, with a focus on Ceratobasicaceae, Tulasnellaceae, and Sebacineaceae. We screened taxa using ITS sequencing, and after contaminants were removed we chose 32 taxa for shallow genome sequencing. A total of 50 taxa were analyzed and we extracted 408 orthologous genes. Two highly-supported phylogenetic trees were constructed with RAxML and ASTRAL-III that were overall highly congruent. We discuss how our study provides new insight into the relationships of these orchid mycorrhizal fungi, highlights areas for taxonomic attention and we suggest future research directions.

## 2. MATERIALS AND METHODS

### 2.1 Taxonomic sampling

32 environmental samples were isolated from endangered orchids. These samples span three genera in three families in two orders. Outgroup genomes were chosen from the repository in Mycocosm to capture the breadth of taxonomic diversity (Grigoriev et al., 2014). Two super outgroups (*Kockovaella* sp and *Calocera sp)* were chosen from the successively sister classes outside the ingroup class Agaricomycetes [Tremellomycetes, [Dacrymycetes, [Agaricomycetes]]]. In the Cantharellales we sampled the three genomes in Ceratobasidiaceae (*Rhizoctonia solani*, *Thanatephorus cucumeris*, and *Ceratobasidium sp AG1)*, the two genomes in Tulasnellaceae (*Tulasnella calospora* AL13/4D, and *Tulasnella calospora* UAMH9824), and one genome each from 4 of the remaining 5 families *Botryobasidium botryosum* (Botryobasidiaceae), *Clavulina sp* (Clavulinaceae), *Cantharellus anzutake* (Cantharellaceae), and *Hydnum rufescens* (Hydnaceae). We also included three genomes in Serendipitaceae (Sebacinales) *Sebacina vermifera* (syn. *Serendipita vermifera)*, *Piriformospora indica* (syn. *Serendipita indica*), and *Serendipita sp.* 407. We sampled representatives from the order Auriculariales to capture the entire diversity of these sequences (*Oliveonia pauxilla, Auricularia subglabra, Aporpium caryae, and Exidia glandulosa*).

### 2.2 Fungal Isolates

The 32 fungal samples used in this study were isolated from roots or protocorms (the seedling stage) of endangered orchid species in areas spanning from Hawaii to Florida, with a focus on the Midwest and the Florida Panther National Wildlife Refuge (Table 1). For the full description of the isolation techniques used, see Zettler and Corey (2018). Briefly, root tissue was surface-sterilized then placed in a petri dish with sterile water and finely diced with a scalpel. Fungal Isolation Media (Clements et al., 1986) was poured on the diced root tissue and left at ambient temperature. After 24-48 hours, the plates were examined with a dissecting microscope to identify fungal growth. Mycelia were excised and placed on Difco Potato Dextrose Agar (PDA; Becton Dickinson and Co., Sparks, MD, Mfr # BD 213400). Those fungi with morphological characteristics consistent with fungi in the form genus *Rhizoctonia* as identified in Currah et al., (1997) were retained for identification with ITS sequencing (Figure 1).

**Table 1.** Description of fungal isolates. UAMH numbers refer to the repository number for isolates deposited in the UAMH Centre for Global Microfungal Diversity.

| Sample ID | Species | Strain | Orchid source | Tissue source | Location |
| --- | --- | --- | --- | --- | --- |
| Cerato11750 | Ceratobasidium sp | UAMH11750 | *Dendrophylax lindenii* (Lindl.) Benth. ex Rolfe | root | Florida Panther National Wildlife Refuge (NWR) |
| Cerato370 | Ceratobasidium sp | 370 | *Platanthera leucophaea* (Nutt.) Lindl. | root | Tuscola Co., MI |
| Cerato392 | Ceratobasidium sp | 392 | *Campylocentrum pachyrrhizum* (Rchb.f.) Rolfe | root | Florida Panther National Wildlife Refuge (NWR) |
| Cerato394 | Ceratobasidium sp | 394 | *Dendrophylax lindenii* (Lindl.) Benth. ex Rolfe | root | Florida Panther National Wildlife Refuge (NWR) |
| Cerato395 | Ceratobasidium sp | 395 | *Campylocentrum pachyrrhizum* (Rchb.f.) Rolfe | root | Florida Panther National Wildlife Refuge (NWR) |
| Cerato414 | Ceratobasidium sp | 414 | *Platanthera lacera* (Michx.) G.Don | root | Fayette Co., IL |
| Cerato423 | Ceratobasidium sp | 423 | *Spiranthes vernalis* Engelm. & A.Gray | root | Madison Co., IL |
| Cerato428 | Ceratobasidium sp | 428 | *Dendrophylax porrectus* (Rchb.f.) Carlsward & Whitten | root | Florida Panther National Wildlife Refuge (NWR) |
| Serend396 | Serendipita sp | 396 | *Prosthechea cochleata* (L.) W.E.Higgins | root | Florida Panther National Wildlife Refuge (NWR) |
| Serend397 | Serendipita sp | 397 | *Prosthechea cochleata* (L.) W.E.Higgins | root | Florida Panther National Wildlife Refuge (NWR) |
| Serend398 | Serendipita sp | 398 | *Prosthechea cochleata* (L.) W.E.Higgins | root | Florida Panther National Wildlife Refuge (NWR) |
| Serend399 | Serendipita sp | 399 | *Encyclia tampensis* Small | root | Florida Panther National Wildlife Refuge (NWR) |
| Serend400 | Serendipita sp | 400 | *Encyclia tampensis* Small | root | Florida Panther National Wildlife Refuge (NWR) |
| Serend401 | Serendipita sp | 401 | *Epidendrum amphistomum* A.Rich | root | Florida Panther National Wildlife Refuge (NWR) |
| Serend405 | Serendipita sp | 405 | *Prosthechea cochleata* (L.) W.E.Higgins | root | Florida Panther National Wildlife Refuge (NWR) |
| Serend407 | Serendipita sp | 407 | *Epidendrum amphistomum* A.Rich | root | Florida Panther National Wildlife Refuge (NWR) |
| Serend411 | Serendipita sp | 411 | *Prosthechea cochleata* (L.) W.E.Higgins | root | Florida Panther National Wildlife Refuge (NWR) |
| Tulasn330 | Tulasnella sp | 330 | *Platanthera holochila* (Hillebr.) Kraenzl. | peloton | Molokai, HI |
| Tulasn331 | Tulasnella sp | 331 | *Platanthera holochila* (Hillebr.) Kraenzl. | peloton | Molokai, HI |
| Tulasn332 | Tulasnella sp | 332 | *Platanthera holochila* (Hillebr.) Kraenzl. | peloton | Molokai, HI |
| Tulasn403 | Tulasnella sp | 403 | *Oeceoclades maculata* Lindl. | root | Florida Panther National Wildlife Refuge (NWR) |
| Tulasn408 | Tulasnella sp | 408 | *Polystachya concreta* (Jacq.) Garay & H.R.Sweet | root | Florida Panther National Wildlife Refuge (NWR) |
| Tulasn417 | Tulasnella sp | 417 | *Platanthera leucophaea* (Nutt.) Lindl. | root | McHenry Co., IL |
| Tulasn418 | Tulasnella sp | 418 | *Platanthera leucophaea* (Nutt.) Lindl. | root | McHenry Co., IL |
| Tulasn419 | Tulasnella sp | 419 | *Cypripedium candidum* Muhl. ex Willd. | protocorm/seedling | McHenry Co., IL |
| Tulasn424 | Tulasnella sp | 424 | *Platanthera paramoena* A.Gray | root | Fayette Co., IL |
| Tulasn425 | Tulasnella sp | 425 | *Platanthera paramoena* A.Gray | root | Fayette Co., IL |
| Tulasn427 | Tulasnella sp | 427 | *Dendrophylax porrectus* (Rchb.f.) Carlsward & Whitten | root | Florida Panther National Wildlife Refuge (NWR) |
| Tulasn9824 | Tulasnella calospora | UAMH9824 | *Spiranthes brevilabris* Lindl. | root | Levy Co., FL |
| Tulinq235 | Tulasnella inquilina | 235 | *Platanthera integrilabia* (Correll) Luer | root | McMinn Co., TN |
| Tulinq238 | Tulasnella inquilina | 238 | *Platanthera integrilabia* (Correll) Luer | root | McMinn Co., TN |
| Tulinq7632 | Tulasnella inquilina | UAMH7632 | *Platanthera integrilabia* (Correll) Luer | root | Greenville, SC |

**Figure 1.**
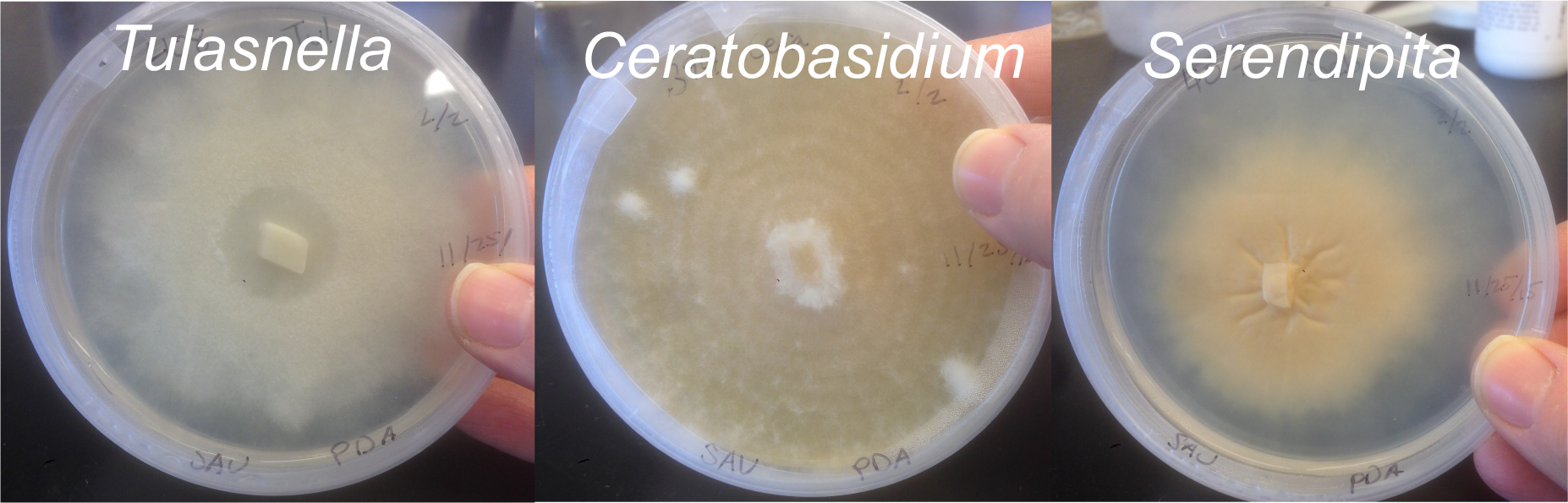
Morphological examples of Tulasnella, Ceratobasidium, and Serendipita. One representative from each genus from the Zettler collection. All three isolates started growing on Potato Dextrose Agar on the same day as indicated by the date on the petri dish (25 November 2015). Photographs: Sarah Unruh.

Fungi were grown in flasks with 75ml of full strength Difco Potato Dextrose Broth (Difco Becton Dickinson and Co., Sparks, MD, Mfr # BD 254920) on a shaker table until there was enough tissue for extraction. Depending on the isolate this took 2-6 weeks. Often multiple flasks of each isolate were grown at one time to speed up this process. For extraction, the entire contents of each flask was poured into a 150mL Polystyrene Bottle Top Filter 0.45um (Corning Incorporated, Corning, USA, Cat # 430627) and washed with DI water. These samples were weighed to determine how many samples could be processed from each sample (minimum of 0.2 grams filtered weight/tube). Fungi were isolated with either the Bacterial/Fungal DNA extraction kit (Zymo Research, Irvine, USA, Cat # D6005, Lot # ZRC201856) according to manufacturer protocol or a CTAB, phenol chloroform isoamyl procedure (Supplemental Figure S1). When the Zymo kit was used, fungi were added to lysis tubes and put on bead beater for two rounds of four minutes. If the CTAB extraction was employed, fungal tissue was ground with liquid Nitrogen in ceramic mortar and pestle. Extracted DNA was assayed on a NanoDrop 2000 (ThermoFisher Scientific, USA, cat # ND-2000) and on a Qubit 2.0 Fluorometer (ThermoFisher Scientific, USA, cat # Q32866) with the Qubit double-stranded DNA High Sensitivity Assay kit (ThermoFisher Scientific, USA, cat # Q32851). We followed JGI instruction for sample submission by submitting approximately 500 ng of each sample in a total volume of 25-35 uL in one 96-well plate provided by JGI.

### 2.3 ITS sequencing for Species Identification

To determine species identity, we sequenced the internal transcribed spacer (ITS) region of the rDNA. We used the same DNA extraction methods referenced above. We used the primer pairs ITS1/ITS4-Tul or ITS1-OF/ITS4-OF for isolates presumed to be *Tulasnella* as the ITS sequences in this genus are highly divergent and not captured well with other primers (Taylor and McCormick, 2008). For the genera *Ceratobasidium* and *Serendipita,* the general primers ITS1/ITS4 or ITS1-OF/ITS4-OF were used and if these did not successfully amplify the ITS region of *Serendipita* isolates the primer pair ITS3Seb/NL4 (Bellemain et al., 2010; Ray et al., 2015; White et al., 1990). The amplified DNA was cleaned with the DNA Clean and Concentrator-25 kit (Zymo Research, Irvine, USA, cat # D4033). These PCR products were assessed on a 1.5% agarose gel and Sanger sequencing was performed at the University of Missouri DNA Core Facility. These sequences were evaluated for confidence in base calling and edited by trimming low quality bases from the beginning and end of each sequence in Geneious 9.1.8 (http://www.geneious.com/). These trimmed sequences were queried against NCBI’s default nucleotide-nucleotide database as well as the UNITE database for species identification (Nilsson et al., 2019). These sequences were generated for the purpose of accurate species ID before sending DNA samples for shallow genome sequencing.

### 2.4 Shallow Genome Sequencing and Quality Control

Shallow genome sequencing of 32 samples, quality control, and filtering were performed at the Joint Genome Institute (JGI) under a Community Sequencing Proposal (#2000). Samples were run on an Illumina NovaSeq with 2×151 base pair (bp) reads. The quality control and filtering at the JGI use BBmap to remove contamination and remove low quality reads (Bushnell B., BBMap. http://sourceforge.net/projects/bbmap/). Three samples were sequenced at the University of Missouri’s DNA Core Facility which were run on an Illumina NextSeq 500 machine on one lane with 45 other samples generating 2×150 bp reads.

### 2.5 Shallow Genome Assembly and Annotation

All cleaned and filtered sequences from the Joint Genome Institute and the University of Missouri were assembled with the AAFTF pipeline for read assembly, remove vector contamination and duplicate contigs, contig sequence polishing and sorting the contigs by length (Stajich, JE., Automatic Assembly For the Fungi. https://github.com/stajichlab/AAFTF). The pipeline performs assembly with Spades 3.10.0 using default parameters which consider 3 kmer values and select the optimal assembly based on summary statistics (Nurk et al., 2013). As a measure to assess genome completeness, all samples were run through BUSCO 3.0.2 using the Basidiomycota database (Simao et al., 2015). For most samples, RNA sequence data was used to facilitate annotation. When samples were too distantly related to map efficiently to the RNA sequencing reads, these taxa were annotated without aligning to the RNA sequences (Table 5). The RNA sequences used for reference were also generated from JGI CSP #2000 and will be published as part of a separate study.

All samples were then prepared for gene prediction using Funannotate 1.6.0 (Palmer JP, Stajich JE. 2018, https://github.com/nextgenusfs/funannotate), which performs all the steps necessary for genome annotation from gene prediction training to final gene consensus model, functional prediction, and dataset preparation for deposition into GenBank. The tool first runs RepeatMasker 4.0.7 (http://www.repeatmasker.org). This “softmasks” the genome by converting repetitive elements into lowercase letters in the assembly files. This step is necessary for the gene prediction steps that follow. After masking, each assembly is run through a training step to provide the initial models for the *ab initio* gene prediction programs AUGUSTUS 3.3.0 (Keller et al., 2011; Stanke and Waack, 2003), SNAP (Korf, 2004), CodingQuarry (Testa et al., 2015), and GeneMark-ES/ET 4.38.0 (Lomsadze et al., 2014). Protein sequences are also aligned with diamond (Buchfink et al., 2015) and gene models polished with exonerate (Slater and Birney, 2005). When RNASeq reads were available for a strain, these were applied as part of a training step which first aligned short RNASeq reads, followed by assembly of these reads into contig with Trinity. Finally these assembled transcripts were aligned to the genome to produce gene models which were used for gene predictor training. Table 5 has the strains which were able to use the RNASeq data as support for gene model training and prediction. These combined evidence of these gene predictions, both *ab initio* and protein and transcript sequence based, were combined with EvidenceModeler to use combined evidence to predict a final set of protein coding genes. In addition tRNA gene predictions were performed with tRNAScan-SE (Lowe and Eddy, 1997). The resulting predicted protein files were then used for the phylogenetic analyses.

### 2.6 Phylogenomic analysis

We used the pipeline PHYling 1.0 (https://doi.org/10.5281/zenodo.1257001) developed by the Stajich lab, to extract orthologous genes from the predicted proteins of our taxa (Spatafora et al., 2016). PHYling uses Hmmer3 (v3.2.1) to compare our predicted proteins to a list of Profile-Hidden-Markov models of phylogenetically informative markers. The list we used is the 434 orthologous gene set (https://doi.org/10.5281/zenodo.1251476) constructed by the 1000 Fungal Genomes Project and identified as single-copy in orthologous gene clusters available from the Joint Genome Institute’s MycoCosm repository (Grigoriev et al., 2014). We used hmmsearch to compare each sample’s proteome to the 434 gene list. The protein sequence homologs we identified were aligned to the marker-profile HMM with hmmalign. These alignments were concatenated to run a phylogenetic analysis with RAxML 8.2.12 (Stamatakis, 2006; Stamatakis et al., 2008). The model of evolution was determined automatically and bootstrapped with 100 replicates. The gene trees generated from RAxML were used to construct a consensus tree with ASTRAL-III 5.6.3 (Mirarab et al., 2014; Zhang et al., 2017).

### 2.7 Data accessibility

Isolates with UAMH numbers are stored in the UAMH Centre for Global Microfungal Biodiversity repository. Raw DNA sequence data have been deposited in SRA and are associated with BioProjects listed in Table 3. Scripts used for these analyses and all alignments, trees, and intermediate files will be made available in a Dryad repository upon publication. BioProject IDs and JGI Mycocosm repositories are summarized in Table 3.

## 3. RESULTS

### 3.1 ITS identifications

For the 35 isolates studied, ITS identifications, primers used and the length of each sequence are summarized in Table 2. One sample sent to the Joint Genome Institute was not sequenced due to poor DNA quality. Two isolates were identified as contaminants (isolates 420 and 422) and were excluded from further analysis (Table 2). Only four out of 35 isolates were identified to species.

**Table 2.** Identifications of fungal isolates based on the internal transcribed spacer (ITS)

| SampleID | Morphological ID | Top hit UNITE | Top hit GenBank | Primer sequenced | Length edited in base pairs |
| --- | --- | --- | --- | --- | --- |
| Cerato11750 | *Ceratobasidium* | Ceratobasidiaceae | Uncultured *Ceratobasidium* clone LP8-Cer1 | ITS1 | 641 |
| Cerato370 | *Ceratobasidium* | *Ceratobasidium* | *Ceratobasidium* UAMH 9847 | ITS4 | 538 |
| Cerato392 | *Ceratobasidium* | Basidiomycota (same as ncbi) | orchid mycorrhizae KH4-8 | ITS4 | 550 |
| Cerato394 | *Ceratobasidium* | *Ceratobasidium* | *Ceratobasidium* | ITS1 | 586 |
| Cerato395 | *Ceratobasidium* | Ceratobasidiaceae | *Ceratobasidium* sp JTO161 | ITS1 | 548 |
| Cerato414 | *Ceratobasidium* | Ceratobasidiaceae | *Ceratobasidium* sp | ITS1 | 100 |
| Cerato423 | *Ceratobasidium* | *Ceratobasidium* | Uncultured Ceratobasidiaceae clone 207 | ITS4 | 390 |
| Cerato428 | *Ceratobasidium* | Ceratobasidiaceae | *Ceratobasidium* sp JTO161 | ITS1 | 420 |
| Serend396 | *Serendipita* | Sebacinales (orchid fungus) | uncultured Sebacinales clone | ITS1 | 189 |
| Serend397 | *Serendipita* | Sebacinales | uncultured Sebacinales clone | NL4 | 680 |
| Serend398 | *Serendipita* | Sebacinales | uncultured Sebacinales clone | NL4 | 567 |
| Serend399 | *Serendipita* | Sebacinales | uncultured Sebacinales clone | NL4 | 740 |
| Serend400 | *Serendipita* | Sebacinales | *Serendipita* sp MAFF 305831 | NL4 | 780 |
| Serend401 | *Serendipita* | Sebacinales | Uncultured Sebacinales clone LP49-23S | ITS4 | 614 |
| Serend405 | *Serendipita* | *Serendipita* | *Serendipita* sp MAFF 305831 | NL4 | 380 |
| Serend407 | *Serendipita* | Sebacinales | Uncultured *Sebacina* mycobiont of *Riccardia palmata* | * | 2316 |
| Serend411 | *Serendipita* | Sebacinales | Uncultured Sebacinales clone LP49-23S | ITS3Seb | 880 |
| Tulasn330 | *Tulasnella* | Tulasnellaceae | Uncultured Tulasnellaceae isolate 55P-Leu13 | ITS4 | 570 |
| Tulasn331 | *Tulasnella* | Tulasnellaceae | Uncultured Tulasnellaceae | ITS1 | 620 |
| Tulasn332 | *Tulasnella* | Tulasnellaceae | Uncultured Tulasnellaceae | ITS4-OF | 810 |
| Tulasn403 | *Tulasnella* | *Tulasnella* | *Tulasnella* sp CH01 | ITS1 | 490 |
| Tulasn408 | *Tulasnella* | Tulasnellaceae | Uncultured Tulasnellaceae clone DOF-YC9 | ITS4 | 622 |
| Tulasn417 | *Tulasnella* | *Tulasnella* | *Tulasnella* sp 9 MM-2012 | ITS1 | 870 |
| Tulasn418 | *Tulasnella* | Tulasnellaceae | Uncultured Tulasnellaceae P94 | ITS1 | 350 |
| Tulasn419 | *Tulasnella* | *Tulasnella* | Tulasnellaceae sp Pch 253 | ITS4 | 368 |
| Tulasn424 | *Tulasnella* | Tulasnellaceae | Tulasnellaceae | ITS4 | 605 |
| Tulasn425 | *Tulasnella* | *Tulasnella* | *Tulasnella* sp 149 | ITS1 | 570 |
| Tulasn427 | *Tulasnella* | Tulasnellaceae | Uncultured *Tulasnella* clone 998OF | ITS4 | 380 |
| Tulasn9824 | *Tulasnella calospora* | *Tulasnella calospora* | *Tulasnella calospora* isolate Pch-QS-0-1 | ITS4-Tul | 148 |
| Tuling235 | *Epulorhiza inquilina* | *Tulasnella* | *Tulasnella* sp 3MV-2011 PA 053A | ITS4 | 650 |
| Tuling238 | *Epulorhiza inquilina* | Tulasnellaceae | *Tulasnella* sp 3MV-2011 PA 053A | ITS4 | 758 |
| Tuling7632 | *Epulorhiza inquilina* | *Tulasnella* | *Tulasnella* sp 3MV-2011 PA 053A | ITS1 | 790 |
| Isolate 420* | *Tulasnella* | *Phanerochaete australis* | *Phanerochaete australis* | ITS1 | 350 |
| Isolate 422* | *Tulasnella* | *Trichoderma petersenii* | *Trichoderma* sp isolate ARMI-23 | ITS4 | 390 |

**Table 3.** List of taxa and data availability.

| **SampleID** | **BioProject or JGI web portal** | **BioSample** |
| --- | --- | --- |
| * *Ceratobasidium_sp_UAMH11750*.Orchid | PRJNA558776 | SAMN12498506 |
| *Ceratobasidium_sp_370*.Orchid | PRJNA557749 | SAMN12427929 |
| *Ceratobasidium_sp_392*.Orchid | PRJNA557750 | SAMN12427914 |
| * *Ceratobasidium_sp_394*.Orchid | PRJNA557751 | SAMN12427897 |
| * *Ceratobasidium_sp_395*.Orchid | PRJNA557752 | SAMN12427926 |
| *Ceratobasidium_sp_414*.Orchid | PRJNA557753 | SAMN12427925 |
| * *Ceratobasidium_sp_423*.Orchid | PRJNA557754 | SAMN12427910 |
| *Ceratobasidium_sp_428*.Orchid | PRJNA557755 | SAMN12427923 |
| *Serendipita_sp_396*.Orchid | PRJNA557757 | SAMN12427894 |
| *Serendipita_sp_397*.Orchid | PRJNA557758 | SAMN12427928 |
| *Serendipita_sp_398*.Orchid | PRJNA557759 | SAMN12427900 |
| *Serendipita_sp_399*.Orchid | PRJNA557760 | SAMN12427906 |
| * *Serendipita_sp_400*.Orchid | PRJNA557761 | SAMN12427895 |
| *Serendipita_sp_401*.Orchid | PRJNA557762 | SAMN12427903 |
| * *Serendipita_sp_405*.Orchid | PRJNA557763 | SAMN12427917 |
| * *Serendipita_sp_407*.Orchid | PRJNA558790 | SAMN12498938 |
| * *Serendipita_sp_411*.Orchid | PRJNA557734 | SAMN12427911 |
| *Tulasnella_sp_330*.Orchid | PRJNA557739 | SAMN12427924 |
| *Tulasnella_sp_331*.Orchid | PRJNA557740 | SAMN12427908 |
| * *Tulasnella_sp_332*.Orchid | PRJNA557741 | SAMN12427902 |
| *Tulasnella_sp_403*.Orchid | PRJNA557742 | SAMN12427920 |
| *Tulasnella_sp_408*.Orchid | PRJNA557743 | SAMN12427916 |
| *Tulasnella_sp_417*.Orchid | PRJNA557744 | SAMN12427919 |
| *Tulasnella_sp_418*.Orchid | PRJNA557745 | SAMN12427921 |
| * *Tulasnella_sp_419*.Orchid | PRJNA557746 | SAMN12427922 |
| *Tulasnella_sp_424*.Orchid | PRJNA557747 | SAMN12427899 |
| * *Tulasnella_sp_425*.Orchid | PRJNA557748 | SAMN12427912 |
| * *Tulasnella_sp_427*.Orchid | PRJNA557733 | SAMN12427904 |
| * *Tulasnella_calospora_UAMH9824*.Orchid | PRJNA558788 | SAMN12498837 |
| *Tulasnella_inquilina_235*.Orchid | PRJNA557736 | SAMN12427891 |
| * *Tulasnella_inquilina* _238.Orchid | PRJNA557737 | SAMN12427893 |
| * *Tulasnella_inquilina* _UAMH7632.Orchid | PRJNA557738 | SAMN12427890 |
| *Aporpium_caryae*_L-13461. | https://mycocosm.jgi.doe.gov/Elmca1 |  |
| *Auricularia_subglabra*_v2.0 | https://mycocosm.jgi.doe.gov/Aurde3_1 |  |
| *Botryobasidium_botryosum*_v1.0 | https://mycocosm.jgi.doe.gov/Botbo1 |  |
| *Calocera_cornea*_v1.0 | https://mycocosm.jgi.doe.gov/Calco1 |  |
| *Cantharellus_anzutake*_C23_v1.0 | https://mycocosm.jgi.doe.gov/Cananz1 |  |
| *Clavulina_sp._PMI_390_v1.0* | https://mycocosm.jgi.doe.gov/ClaPMI390 |  |
| *Exidia_glandulosa*_v1.0 | https://mycocosm.jgi.doe.gov/Exigl1 |  |
| *Hydnum_rufescens*_UP504_v2.0 | https://mycocosm.jgi.doe.gov/Hydru2 |  |
| *Kockovaella_imperatae*_NRRL_Y-17943_v1.0 | https://mycocosm.jgi.doe.gov/Kocim1 |  |
| *Oliveonia_pauxilla*_MPI-PUGE-AT-0066_v1.0 | https://mycocosm.jgi.doe.gov/Olipa1 |  |
| *Piriformospora_indica*_DSM_11827_from_MPI | https://mycocosm.jgi.doe.gov/Pirin1 |  |
| *Rhizoctonia_solani*_AG-1_IB | https://mycocosm.jgi.doe.gov/Rhiso1 |  |
| *Sebacina_vermifera*_MAFF_305830_v1.0 | https://mycocosm.jgi.doe.gov/Sebve1 |  |
| *Serendipita_sp._407_v1.0* | https://mycocosm.jgi.doe.gov/Serend1 |  |
| *Thanatephorus_cucumeris*_MPI-SDFR-AT-0096_v1.0 | https://mycocosm.jgi.doe.gov/Thacu1 |  |
| *Tulasnella_calospora*_AL13_4D_v1.0 | https://mycocosm.jgi.doe.gov/Tulca1 |  |

Asterisks \* indicate isolates selected for reference genome sequencing.

### 3.2 Shallow genome sequencing and annotation

Shallow genome sequencing of 32 fungal isolates resulted in a wide range in the number of genes annotated in each individual genome. The isolate *Serendipita sp* 396 has the least number of annotated genes at 8,285 and *Ceratobasidium sp* 428 has the most at 25,099. The BUSCO completeness scores ranged from 54.2% to 96.6% of the 1335 orthologues in the BUSCO dataset. For assembly statistics see Table 4 and for BUSCO completeness scores see Table 5. Out of 435 orthologous genes, 429 had enough significant hits for further analysis. The number of genes present for each taxa ranged from 299 in *Tulasnella sp* 408 to 425 in the outgroups *Auricularia subglabra* and *Botryobasidium botryosum*. For full matrix occupancy see Table 6. The outgroup *Kockovaella imperatae* contained 408 of the 429 genes so those 408 sequences were included in the phylogenetic analyses. The concatenated alignment has 128,774 distinct alignment patterns and is 14.31% gaps.

**Table 4.** Assembly statistics.

| SampleID | CONTIG<br>COUNT | TOTAL LENGTH | MIN | MAX | MEDIAN | MEAN | L50 | N50 | L90 | N90 |
| --- | --- | --- | --- | --- | --- | --- | --- | --- | --- | --- |
| Ceratobasidium_sp_UAMH11750.Orchid | 7239 | 47766782 | 2000 | 131072 | 4320 | 6598.53 | 1452 | 8784 | 5266 | 2947 |
| Ceratobasidium_sp_370.Orchid | 8028 | 47284605 | 2000 | 145775 | 3838 | 5889.96 | 1613 | 7304 | 5991 | 2715 |
| Ceratobasidium_sp_392.Orchid | 8904 | 52465834 | 1500 | 100768 | 3806 | 5892.39 | 1693 | 8316 | 6228 | 2516 |
| Ceratobasidium_sp_394.Orchid | 8562 | 53454716 | 1500 | 94250 | 3864 | 6243.25 | 1547 | 9440 | 5859 | 2601 |
| Ceratobasidium_sp_395.Orchid | 8769 | 67010313 | 1500 | 125675 | 4575 | 7641.73 | 1478 | 12298 | 5716 | 3127 |
| Ceratobasidium_sp_414.Orchid | 4161 | 50425407 | 1500 | 342430 | 4339 | 12118.58 | 349 | 34639 | 2143 | 4179 |
| Ceratobasidium_sp_423.Orchid | 15380 | 66434938 | 1500 | 107431 | 2946 | 4319.57 | 3306 | 5259 | 11578 | 2042 |
| Ceratobasidium_sp_428.Orchid | 8999 | 69097995 | 1500 | 121965 | 4552 | 7678.41 | 1493 | 12317 | 5876 | 3139 |
| Serendipita_sp_396.Orchid | 1431 | 20638744 | 2072 | 267195 | 9279 | 14422.6 | 279 | 17663 | 1083 | 6736 |
| Serendipita_sp_397.Orchid | 4253 | 28775535 | 1500 | 270096 | 3582 | 6765.94 | 580 | 10829 | 2793 | 2618 |
| Serendipita_sp_398.Orchid | 4302 | 28851264 | 1500 | 267525 | 3546 | 6706.48 | 574 | 11054 | 2835 | 2583 |
| Serendipita_sp_399.Orchid | 5013 | 31004825 | 1500 | 127831 | 3844 | 6184.88 | 906 | 8927 | 3450 | 2622 |
| Serendipita_sp_400.Orchid | 4392 | 28560853 | 1502 | 143991 | 3564 | 6502.93 | 635 | 10584 | 2927 | 2562 |
| Serendipita_sp_401.Orchid | 3823 | 28571286 | 1500 | 662216 | 3626 | 7473.52 | 429 | 13497 | 2433 | 2804 |
| Serendipita_sp_405.Orchid | 4254 | 28724145 | 1500 | 280457 | 3539 | 6752.27 | 566 | 10892 | 2796 | 2594 |
| Serendipita_sp_407.Orchid | 4028 | 27230538 | 1500 | 297010 | 3154 | 6760.31 | 426 | 12923 | 2590 | 2442 |
| Serendipita_sp_411.Orchid | 4211 | 28400685 | 1500 | 296955 | 3574 | 6744.4 | 574 | 10730 | 2767 | 2605 |
| Tulasnella_sp_330.Orchid | 1013 | 42302809 | 1500 | 967722 | 8117 | 41759.93 | 66 | 175815 | 323 | 19296 |
| Tulasnella_sp_331.Orchid | 3446 | 44512311 | 1500 | 298887 | 4965 | 12917.1 | 311 | 35916 | 1764 | 4762 |
| Tulasnella_sp_332.Orchid | 3329 | 44576393 | 1501 | 416704 | 5260 | 13390.33 | 297 | 36313 | 1699 | 5096 |
| Tulasnella_sp_403.Orchid | 2963 | 29964626 | 1502 | 501591 | 4201 | 10112.93 | 303 | 23340 | 1647 | 3550 |
| Tulasnella_sp_408.Orchid | 10591 | 63626598 | 1500 | 92931 | 3595 | 6007.61 | 1850 | 9042 | 7284 | 2490 |
| Tulasnella_sp_417.Orchid | 9866 | 61487333 | 1500 | 224139 | 3694 | 6232.25 | 1709 | 9469 | 6703 | 2528 |
| Tulasnella_sp_418.Orchid | 3880 | 32841821 | 1501 | 268652 | 3496 | 8464.39 | 411 | 18771 | 2277 | 2844 |
| Tulasnella_sp_419.Orchid | 3865 | 33665047 | 1500 | 229676 | 3923 | 8710.23 | 419 | 18466 | 2308 | 3120 |
| Tulasnella_sp_424.Orchid | 781 | 48431701 | 1507 | 1054277 | 13975 | 62012.42 | 71 | 195410 | 275 | 36329 |
| Tulasnella_sp_425.Orchid | 769 | 48399368 | 1507 | 1089567 | 16223 | 62938.06 | 74 | 189857 | 290 | 37534 |
| Tulasnella_sp_427.Orchid | 6018 | 39289859 | 1500 | 134256 | 3802 | 6528.72 | 956 | 9789 | 4051 | 2663 |
| Tulasnella_calospora_UAMH9824.Orchid | 4164 | 49802335 | 1500 | 345740 | 5668 | 11960.21 | 481 | 25557 | 2334 | 4747 |
| Tulasnella_inquilina_235.Orchid | 1742 | 44191844 | 1503 | 570014 | 3948 | 25368.45 | 119 | 102565 | 508 | 11931 |
| Tulasnella_inquilina_238.Orchid | 2874 | 45488573 | 1500 | 444022 | 4355 | 15827.62 | 240 | 54071 | 1242 | 5477 |
| Tulasnella_inquilina_UAMH7632.Orchid | 2898 | 46174977 | 1501 | 520136 | 4010 | 15933.39 | 209 | 56439 | 1211 | 5431 |

**Table 5.**
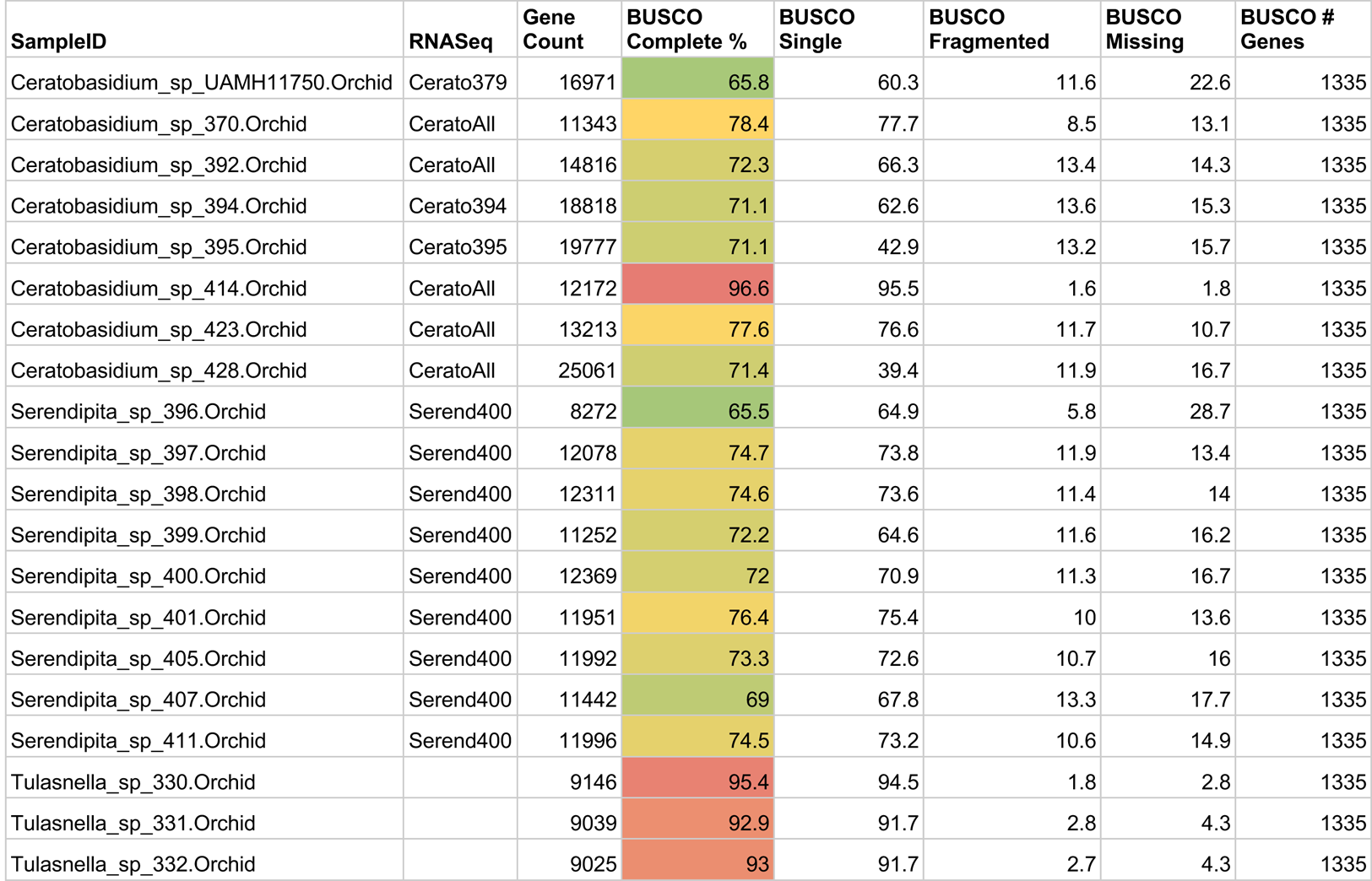
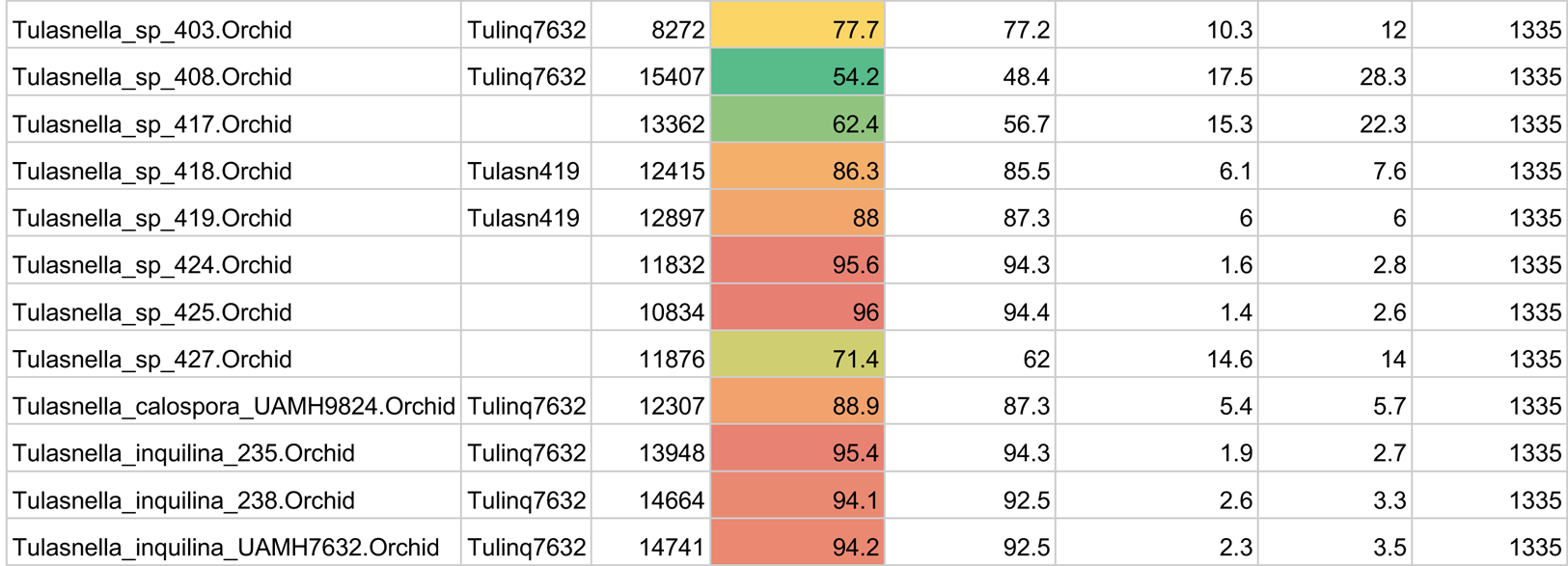
Annotation and BUSCO completeness metrics. Taxa without an RNA sequence listed did not sufficiently map to the Tulinq7632 RNA sequences and were annotated without expression data. The colors in the BUSCO complete % column range from blue-green (lowest percentage) to dark red (highest percentage).

**Table 6.** Matrix Occupany.

| **Sample ID** | **Number best hit genes (429 total)** |
| --- | --- |
| Ceratobasidium_sp_UAMH11750.Orchid | 354 |
| Ceratobasidium_sp_370.Orchid | 356 |
| Ceratobasidium_sp_392.Orchid | 368 |
| Ceratobasidium_sp_394.Orchid | 376 |
| Ceratobasidium_sp_395.Orchid | 369 |
| Ceratobasidium_sp_414.Orchid | 387 |
| Ceratobasidium_sp_423.Orchid | 376 |
| Ceratobasidium_sp_428.Orchid | 382 |
| Serendipita_sp_396.Orchid | 311 |
| Serendipita_sp_397.Orchid | 391 |
| Serendipita_sp_398.Orchid | 371 |
| Serendipita_sp_399.Orchid | 358 |
| Serendipita_sp_400.Orchid | 375 |
| Serendipita_sp_401.Orchid | 382 |
| Serendipita_sp_405.Orchid | 379 |
| Serendipita_sp_407.Orchid | 364 |
| Serendipita_sp_411.Orchid | 382 |
| Tulasnella_sp_330.Orchid | 401 |
| Tulasnella_sp_331.Orchid | 401 |
| Tulasnella_sp_332.Orchid | 398 |
| Tulasnella_sp_403.Orchid | 376 |
| Tulasnella_sp_408.Orchid | 291 |
| Tulasnella_sp_417.Orchid | 330 |
| Tulasnella_sp_418.Orchid | 400 |
| Tulasnella_sp_419.Orchid | 408 |
| Tulasnella_sp_424.Orchid | 408 |
| Tulasnella_sp_425.Orchid | 403 |
| Tulasnella_sp_427.Orchid | 376 |
| Tulasnella_calospora_UAMH9824.Orchid | 398 |
| Tulasnella_inquilina_235.Orchid | 416 |
| Tulasnella_inquilina_238.Orchid | 410 |
| Tulasnella_inquilina_UAMH7632.Orchid | 417 |
| Aporpium_caryae_L-13461. | 423 |
| Auricularia_subglabra_v2.0 | 425 |
| Botryobasidium_botryosum_v1.0 | 425 |
| Calocera_cornea_v1.0 | 410 |
| Cantharellus_anzutake_C23_v1.0 | 412 |
| Ceratobasidium_sp_AGI_v1.0 | 422 |
| Clavulina_sp._PMI_390_v1.0 | 423 |
| Exidia_glandulosa_v1.0 | 422 |
| Hydnum_rufescens_UP504_v2.0 | 413 |
| Kockovaella_imperatae_NRRL_Y-17943_v1.0 | 408 |
| Oliveonia_pauxilla_MPI-PUGE-AT-0066_v1.0 | 419 |
| Piriformospora_indica_DSM_11827_from_MPI | 417 |
| Rhizoctonia_solani_AG-1_IB | 410 |
| Sebacina_vermifera_MAFF_305830_v1.0 | 420 |
| Serendipita_sp._407_v1.0 | 415 |
| Thanatephorus_cucumeris_MPI-SDFR-AT-0096_v1.0 | 424 |
| Tulasnella_calospora_AL13_4D_v1.0 | 405 |
| Tulasnella_calospora_UAMH9824_v1.0 | 426 |

### 3.3 Phylogenetic analysis

The best concatenated tree likelihood is −3406977.36. The bootstrap (BS) support is overall very high with the majority of branches at 100 (Figure 2). Eight branches have bootstrap values below 100, and, of those, only three are below 75. The ASTRAL-III tree shows high congruence with the concatenated tree and all but five branches are supported with 0.7 local posterior probability or higher (Figure 3). The two phylogenies have the exact same topology on the class, order, and family level and recapitulate with high support previously published relationships between orders in the Agaricomycetes [Cantharellales, [Sebacinales, [Auriculariales]]]. The phylogenies are highly congruent within Cantharellales, however, the relationships between *Serendipita* isolates are quite different as discussed below.

**Figure 2.**
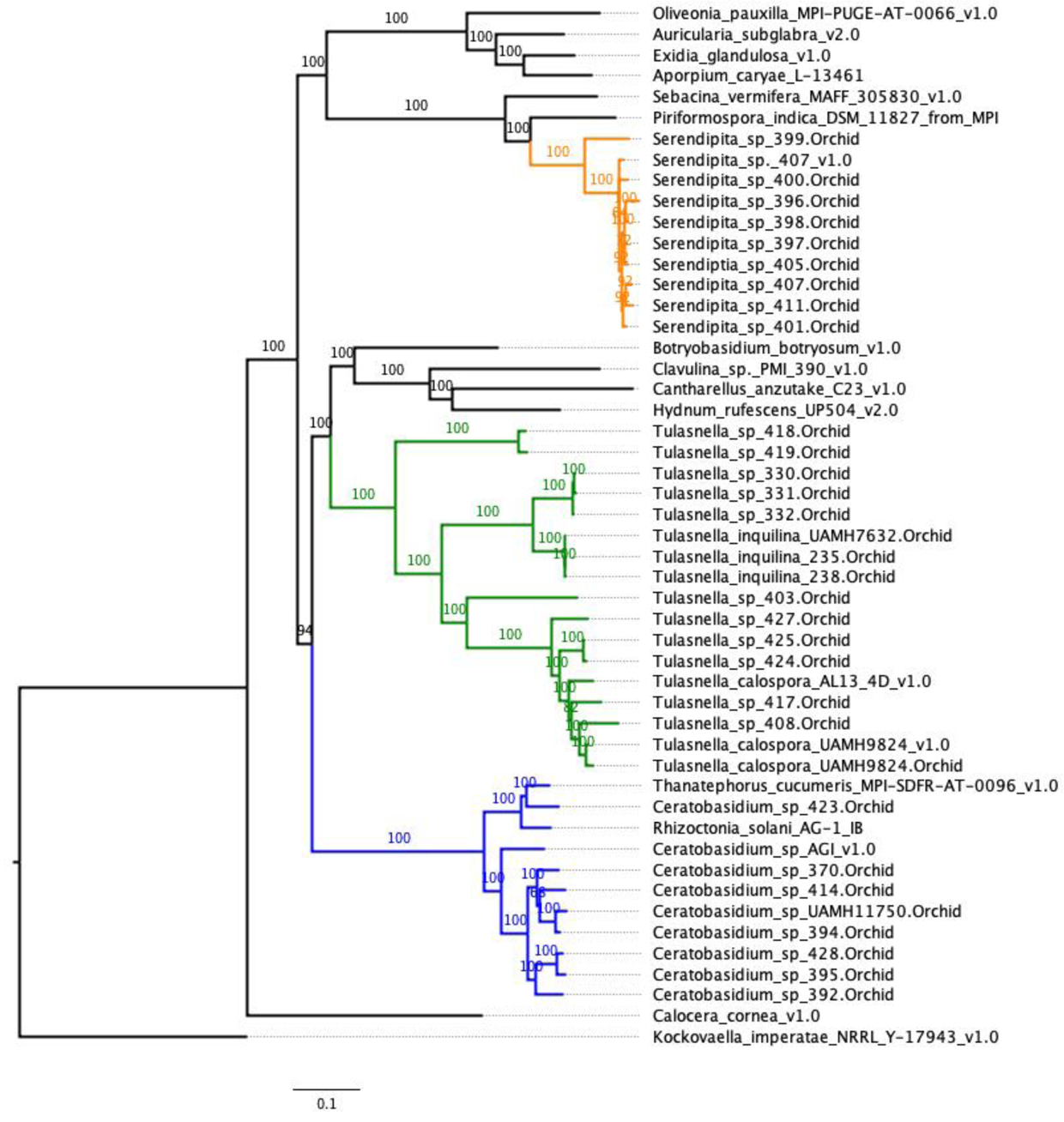
Concatenation-based phylogeny of orchid mycorrhizal fungi. Phylogenetic tree of the orchid mycorrhizal fungi in the Zettler collection with outgroups from the MycoCosm repository (genome.jgi.doe.gov/mycocosm/home). Alignments were made with the Phyling pipeline and the phylogeny was built with RAxML.

**Figure 3.**
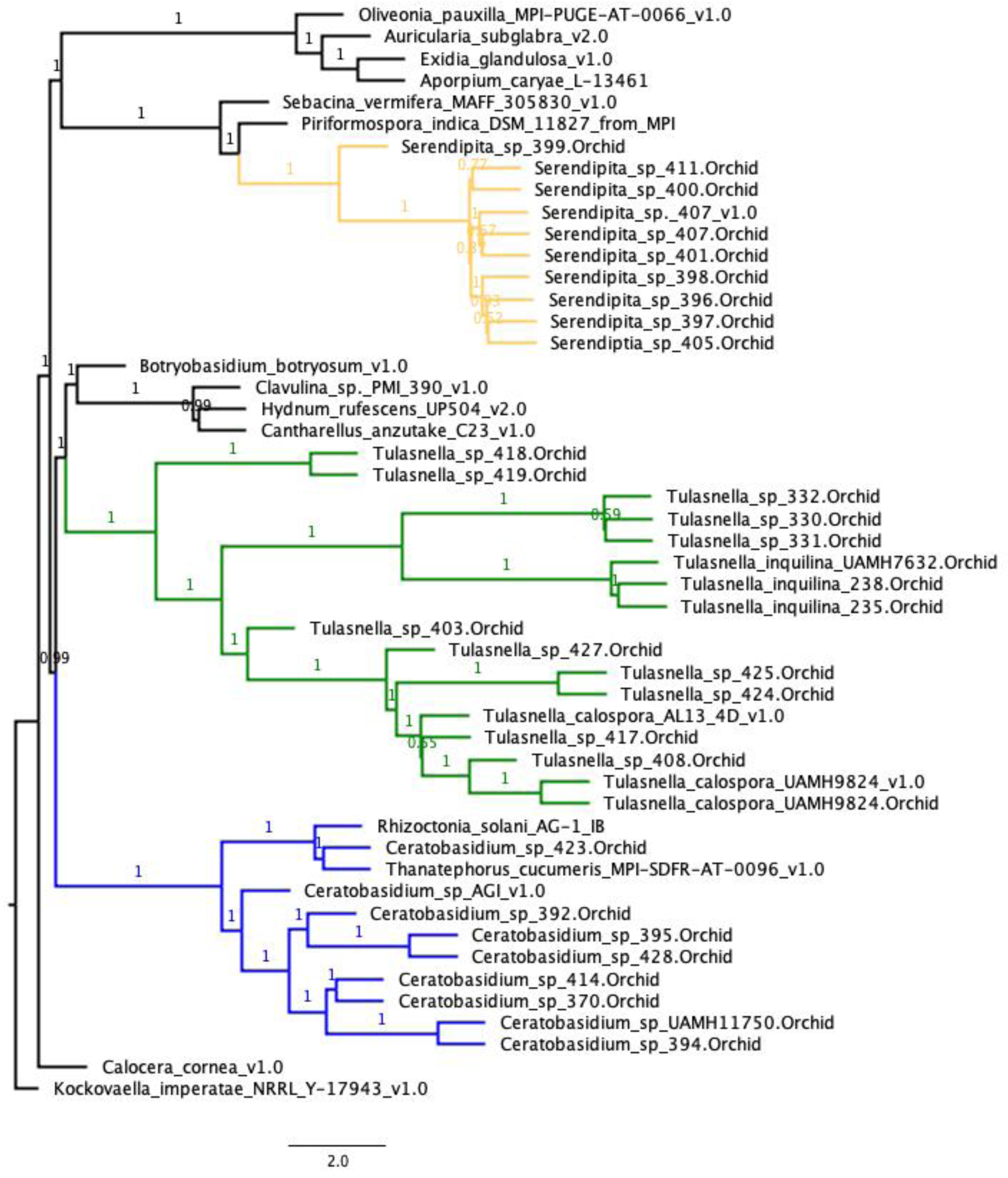
Quartet-based phylogeny of orchid mycorrhizal fungi. Phylogenetic tree of the 32 orchid mycorrhizal fungi in the Zettler collection with 16 outgroups from the MycoCosm repository (genome.jgi.doe.gov/mycocosm/home). Alignments were made with the Phyling pipeline the gene trees were produced with RAxML and the tree was inferred using ASTRAL-III. All posterior probabilities are reported on the tree.

**Figure 4.**
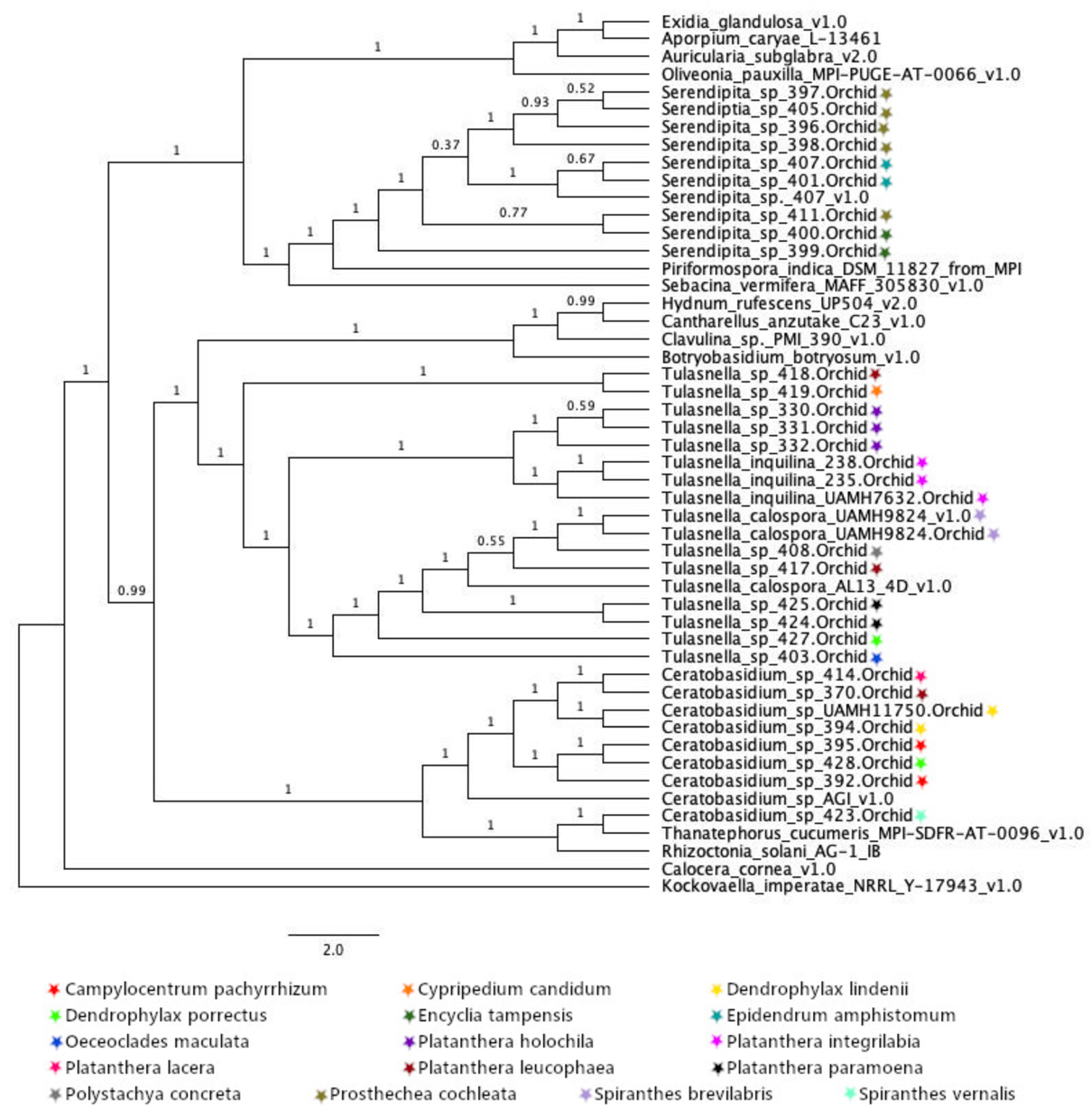
Annotated Quartet-based phylogeny. Phylogenetic tree of the 32 orchid mycorrhizal fungi in the Zettler collection with 18 genomes from the MycoCosm repository (genome.jgi.doe.gov/mycocosm/home). Branches were transformed in FigTree and annotated with colored stars indicating the origin they were isolated from.

Within the Cantharellales, we have strong support (94 BS, 0.99 posterior probability) for Ceratobasidiaceae as sister to the rest of the order. Within Ceratobasidiaceae, the Ceratobasidium isolates cluster together with very strong support with the exception of *Ceratobasidium sp* 423, which is nested within *Rhizoctonia solani* and *Thanatephorus cucumeris*. The only difference between the ML and ASTRAL-III in the family is the placement of *Ceratobasidium sp* 370. In the ML tree, 370 is sister to a clade of [414, [394+UAMH11750]] and in the ASTRAL-III tree, 370 is sister with isolate 414 and equally related to 394+UAMH11750. There is no phylogenetic signal based on orchid source, geographic location (Figure 3, Table 1). Both trees show Tulasnellaceae as sister to the clade [Botryobasidium, [Clavulina, [Cantherellus + Hydnum]]] with 100BS and 1.0 pp. The relationships in *Tulasnella* are highly supported with all but one branch with 100 BS values and all but two branches with pps less than 1.0. Notably, the genome sequence and the shallow genome sequence data for *Tulasnella calospora* UAMH 9824 are sister to each other in the tree, though two other isolates are included in a clade with *Tulasnella calospora* AL13.

The samples in the Sebacinales are not as well-resolved. The *Serendipia* isolates have the least support overall due to the short branches of all isolates aside from *Serendipita* 399, which is sister to the rest. All *Serendipita spp* in this study are most closely related to *Serendipita* (=*Piriformospora) indica* with 100 BS/1.0. It is important to note our inclusion of the reference genome *Serendipita 407* (Serendipita sp._407_v1.0) and a shallow genome sequence of the same isolate (Serendipita_sp_407.Orchid). In our dataset these two samples are not sister to each other. In the quartet-based ASTRAL-III tree, *Serendipita* 400 and 411 are sister to each other with 0.77 posterior probability, whereas in the concatenated tree, the genome of isolate 407 was sister to the rest of the Serendipita isolates aside from 399. The short branches in this group indicate a small number of changes in the alignment in the ML tree and a high degree of discordance in the ASTRAL-III tree. All of the Serendipita isolates are from epiphytic orchids in the Florida Panther National Wildlife Refuge (Figure 3, Table 1). In the ASTRAL tree, Serendipita spp tend to cluster with orchid source compared to the ML tree.

## 4. DISCUSSION

### 4.1 Overview

The primary goal of this study was to use shallow genome sequencing and phylogenetic methods to uncover the evolutionary relationships in a collection of fungal isolates that interact with endangered orchid species. The secondary goal was to leverage current genomic resources to investigate relationships among the orders, families and genera of Agaricomycetes, with a focus on Ceratobasicaceae, Tulasnellaceae, and Sebacineaceae. Understanding of species in the fungal genera that facilitate orchid germination is extremely poor, as the number of formally described species is much lower than the diversity of fungi revealed from metagenomic or environmental sequencing. The results of this study add to our understanding of the genetic diversity of these fungal taxa and provide an example of how sequence data can be incorporated with taxonomic expertise to better describe fungal species.

The fungi that help germinate orchids were first categorized under one “form genus” called *Rhizoctonia* (Currah et al., 1997). This classification is not phylogenetically informative and today we know many orchid symbionts come from two orders (Cantharellales and Sebacinales) in the class Agaricomycetes (Hibbett, 2006). However, the taxonomy remains to be fully resolved. One reason classification can be difficult in these taxa is that these isolates do not sporulate or make sexual structures in laboratory conditions. Another is that traditionally, fungi were classified under two different names – the sexual stage (teleomorph) or vegetative state (anamorph). This policy ended during the 2011 International Botanical Congress when the Nomenclature Section voted to eliminate this dual nomenclature system (Hibbett and Taylor, 2013). Many of the names published in literature are no longer considered the correct taxonomy though in many cases these changes are not strongly reinforced. This study examines the phylogenetic relationships of a collection of isolates so that the genetic distance of these strains is known and to provide a framework for future evolutionary questions. Data from these phylogenies can also provide evidence for new species or to revise current species concepts. Understanding of taxonomy and species relationships is critical for testing evolutionary hypotheses. Increased sampling within taxonomic groups and from sites around the globe is necessary for future studies.

### 4.2 Relationships among Orders and Families

We used shallow genome sequencing for phylogenomics to describe the evolutionary relationships among a collection of orchid mycorrhizal fungi. We also included numerous outgroups to span the amount of biodiversity represented by these fungi. The large number of coding genes allowed us to provide strong evidence for relationships between orders and a novel result within the families of Cantharellales. Our results show strong support for the relationships [Cantharellales, [Sebacinales, [Auriculariales]]]. This is consistent with previously reported studies (Nagy et al,. 2016). Within Cantharellales, the taxonomy is less certain and is still undergoing changes. For example, Dictionary of the Fungi lists seven families while Hibbett et al., (2014) claim four by defining Clavulinaceae and Cantharellaceae as synonymous with Hydnaceae. This decision seems to be based on the authors’ interpretations as the data in the papers they cite don’t support this conclusion (Leacock 2018). Gónzalez et al., (2016), found some support for the relationships [Tulasnellaceae, [Ceratobasidiaceae +Botryobasidiaceae, [Hydnaceae]]] based on the markers ITS-LSU, rpb2, tef1, and atp6. They did state that multiple coding genes would be necessary to see if their result was robust (Gónzalez et al., 2016). Our results show strong support (99 BS and .94 posterior probability) for Ceratobasidiaceae as the sister family to [Tulasnellaceae, [Botryobasidiaceae + rest of Cantharellales]]. We did only include one sample from the four groups besides Ceratobasidiaceae and Tulasnellaceae so more sampling is needed in this group of fungi to produce a robust and consistent phylogenetic inference.

### 4.3 Relationships in Ceratobasidiaceae

The *Ceratobasidium* samples are closely related with the exception of isolate *Ceratobasidium sp* 423 that is nested within *Rhizoctonia solani* and *Thanatephorus cucumeris* (Figures 2, 3). These results are consistent with Veldre et al., (2013), who found that the genera Ceratobasididum and Thanatephorus are polyphyletic. Given the type specimen for Ceratobasidium has since been placed in the Auriculariales, Oberwinkler et al., (2013a) recommended Ceratobasidium should be renamed Rhizoctonia. Given these taxonomic conundrums, attention is needed to make a robust classification system. Something we found affirming was the close relationship of isolates Ceratobasidium 11750 and Ceratobasidium 394. Based on a nearly identical ITS sequence alignment, these isolates were assumed to be very closely related. This result is noteworthy because they have differential abilities to germinate seeds from the endangered Ghost orchid, *Dendrophylax lindenii*. 394 can germinate seeds but 379 does not. More sampling is needed to compare how the isolates included in our study are related to other *Ceratobasidium spp*. that are in defined Anastomosis Groups.

### 4.4 Relationships in Tulasnellaceae

Our *Tulasnella* isolates show a well-supported monophyletic clade in both phylogenetic trees (Figures 2 and 3). Without further targeted sampling, it is premature to delimit species boundaries; however, one species that could use revision is *Tulasnella calospora*. In both the concatenated and coalescent phylogenies, the two *T. calospora* genomes are not sister to each other but include the isolates 408 and 417, which were not identified as *T. calospora* based on the ITS sequence. This result could be a function of the relatively low number of orthologous genes that we recovered from 408 and 417, 291 and 330 out of 434, respectively (Tables 5 and 6). However, others have voiced concern over the species concept (Melissa McCormick, pers. comm.).

Three isolates in this analysis are from the Hawaiian island of Molokai (330, 331, and 332; Table 1). These isolates cluster very closely in both phylogenies and are sister to three isolates of *Tulasnella inquilina*. These isolates turn pink when exposed to light and have highly divergent ITS sequences from the other Tulasnella isolates in this analysis. The strong support for the monophyly of these Hawaiian samples, and their placement in the tree, suggest a potentially new species. With increased sampling, more robust methods to delineate species boundaries such as those used in (Whitehead et al., 2017) and we will have the power to better describe the diversity of orchid mycorrhizal fungi.

### 4.5 Relationships in Sebacinales

All of the *Serenipita* isolates in this analysis are from the Florida National Wildlife Panther Refuge (NWPR) in Florida and they are associated with three different epiphytic orchid species (Table 1). In both phylogenetic analyses, *Serendipita* 399 is sister to the rest of our samples. Growing on PDA, 399 looks morphologically distinct from the other *Serendipita sp* due to a darker orange pigment and a crustose layer on the surface of the agar. This isolate also grows much more slowly than other Serendipita taxa, it would take longer than four weeks for the fungus to grow to the edge of a standard petri dish. For the remaining samples, it could be, that there is one main species or population of *Serendipita* that grows in orchid roots in the NWPR as their relationships are poorly resolved in the RAxML phylogeny and highly incongruent between the two phylogenies. However, in the ASTRAL analysis, the *Serendipita* isolates cluster somewhat closely by the orchid species from which they were isolated though this is not a strong signal (Figure 3). A more thorough and targeted analysis is required to determine the number of distinct populations of these fungi in the Florida National Panther Wildlife Refuge similar to that conducted by Ruibal et al., (2017) to describe the population structure of *Tulasnella prima* in Australia. It would be interesting to survey the fungi growing in the roots of all plant species in the NWPR to determine the genetic diversity of *Serendipita* across the landscape. Such an experiment would show whether orchids are using a narrow distribution of fungi or if the plants are less discerning but the genetic diversity of the fungi is simply very low.

Another result from our analysis shows that these fungal strains are most closely related to *Piriformospora indica*, a known ectomycorrhizal fungus species (Varma et al., 2001). Many fungi in the order Sebacinales are ecologically characterized as ectomycorrhizal fungi and interact with a wide diversity of plant species (Kohler et al., 2015). Indeed, researchers are isolating fungi in the Sebacinales from plants like switchgrass (*Panicum virgatum*) to determine the benefit of these fungi for applications in agriculture (Craven and Ray, 2019). Orchids might contribute to this effort, as it took more than one year for the Craven lab to isolate one strain of *Sebacina vermifera ssp. bescii* from switchgrass; similar fungi are much more easy to isolate from orchid roots (Prasun Ray, pers. comm.). Orchids could be environmental filters for fungi that could be beneficial in many plant-fungal interactions.

### 4.6 Future directions

The next steps stemming from this study are to combine the phylogenetic relationships with taxonomic expertise to name new species or to revisit problematic species concepts like *Tulasnella calospora.* Additionally, it would be beneficial to sequence the genome of the type specimens for many of these genera and species. Being able to compare the genetic sequences of the type specimens would be extremely beneficial for fungal species that do not present sexual characteristics in the lab. A set of fifteen isolates from the collection have been sequenced on the PacBio platform and will be assembled into reference genomes as part of another aim of the Community Sequencing Proposal (Table 3).

## ACKNOWLEDGEMENTS

I thank the Zettler lab students who worked in the field and the lab to isolate these beautiful fungi. We are grateful to the teams of Dr. Greg Bonito, Dr. Daniel Lindner, and Dr. Francis Martin including the ‘Mycorrhizal Genomics Initiative’ consortium and the 1KFG project for access to unpublished genome data. The genome sequence data were produced by the US Department of Energy Joint Genome Institute in collaboration with the user community. The work conducted by the U.S. Department of Energy Joint Genome Institute, a DOE Office of Science User Facility, is supported by the Office of Science of the U.S. Department of Energy under Contract No. DE-AC02-05CH11231. Computations were performed using the computer clusters and data storage resources of the University of California-Riverside HPCC, which were funded by grants from NSF (MRI-1429826) and NIH (1S10OD016290-01A1). Funding for this work is from Joint Genome Institute Community Sequencing Proposal (JGI CSP 2000), National Science Foundation, IOS 1339156, and the Department of Energy (DOE), Defense Threat Reduction Agency (DTRA).

---

**Supplemental Figure S1.**
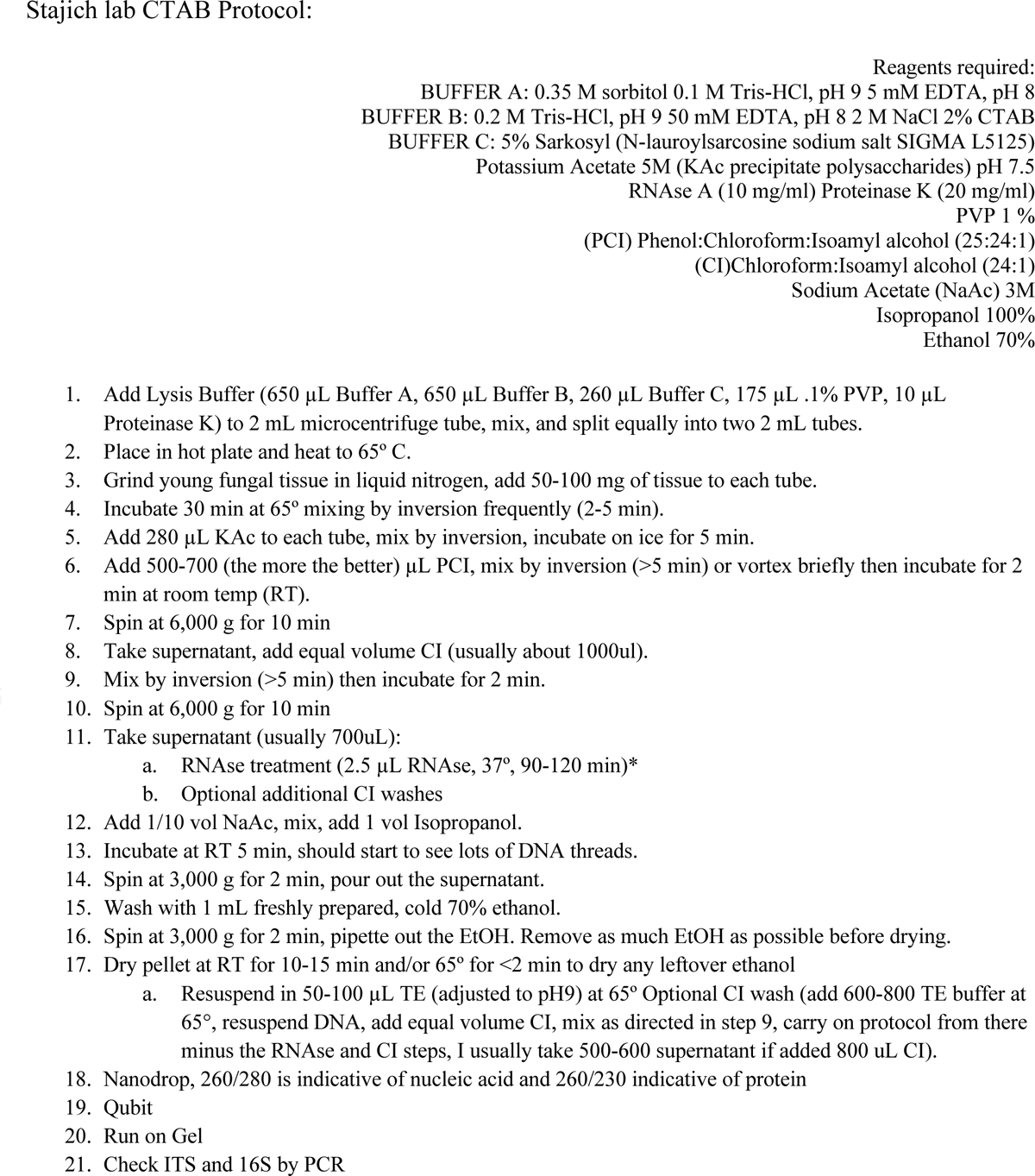
CTAB DNA extraction protocol from Stajich lab.

